# Bayesian analysis of normal mouse cell lineage trees allowing intra-individual, cell population specific mutation rates

**DOI:** 10.1101/040733

**Authors:** Attila Csordas, Remco Bouckaert

## Abstract

Individual cell lineage trees are of biological and clinical importance. The Sanger Mouse Pilot Study used clonal organoid lines of endodermal origin to extract somatic base substitutions from single cells of two healthy mice. Here we applied Bayesian phylogenetics analysis using the Pilot’s 35 somatic base substitutions in order to reconstruct the two cell trees and apply relaxed clock methods allowing intra-individual, cell lineage specific mutation rates. Detailed analysis provided support for the strict clock for mouse_1 and relaxed clock for mouse_2. Interestingly, a clade of two prostate organoid lines in mouse_2 presented one outlier branch mutation rate compared to the mean rates calculated. Based on this study unbiased clock analysis has the potential to add a new, useful layer to our understanding of normal and disturbed cell lineage trees. This study is the first to apply the popular Bayesian BEAST package to individual single cell resolution data.

## Introduction

A fully resolved cell lineage tree of multicellular organisms is a mathematical abstraction, a rooted binary tree that represents all the individual cells, as nodes, and all the divisions, as edges, leading to these cells throughout the life of an individual organism. Understanding even partial cell lineage trees, depicting the lineage relations between a sample of leaf cells from an organism can provide elementary insights into the organisational principles of multicellular biological structures. It might also inform medical decisions in case of diseases, like cancer, where skewed patterns of mutation accumulating cell divisions are introducing imbalances into cell lineage tree structures. In the last decade somatic mutations exposing the genomic variability have been used together with phylogenetic algorithms to reconstruct partial cell lineage trees, primarily in mice using microsatellite instability^1,2^. Somatic base substitutions, a different class of mutations, can also be theoretically used for lineage tree reconstruction besides microsatellites. The increasing use of single cell, whole genome sequencing methods is potentially a big enabler in the field of organismal level cell lineage tree reconstruction^3^. Alternatively, organoid technology deriving clonal lines from individual, organismal leaf cells can be used^4,5^.

The Sanger Mouse Pilot Study (SMPS) generated 25 organoid lines of endodermal origin from stomach (ST), small bowel (SB), large bowel (LB) and prostate (P) of two mice, mouse_1, aged 116 weeks and mouse_2, aged 98 weeks^6^. 35 mutations from these 25 lines have been used for lineage tree topology reconstruction with the help of the maximum parsimony (MP) algorithm.

There are other methods available besides MP. Looking at the microsatellite based cell lineage tree literature, distance matrix - neighbor joining^1^ - and Bayesian methods^2^ were used. From our point of view the most important missing feature of MP reconstruction is that out of the box it provides no help in investigating the level of clock-like behaviour so it does not provide mutation rate estimations of the different lineages.

According to estimations using bulk, whole-exome sequencing of tumour tissues the mutation rate of different normal cell types - lymphocytes, colorectal epithelial cells - is very similar^7^. This suggests the idea that the mutation rates in human cell types are fixed at the same level throughout the body, although the data only covers the minority of all cell types or tissues. Single cell derived data for calculating per division or per unit time mutation rates in mammals (see for instance Wang et al. 2012 studying human germline rates^8^) can be used to investigate whether the same or different cell or tissue populations can exhibit different mutational loads. Single cell derived data has a natural advantage over bulk sequencing approaches when it comes to calculating more fine grained per cell division mutation rates. Since the concept of evolvable mutation rates at every scale is central to our understanding of biology it is a valid question to investigate clock-likeness in case of the single cell derived SMPS data to elaborate the potential mutational dynamics of cell lineage trees. In sync with the literature the default and traditional assumption would be that of a strict molecular clock with a constant substitution rate across all lineages on all sites in an organismal cell lineage tree. One well established class of phylogenetics methods is Bayesian evolutionary analysis. Bayesian phylogenetics tries to estimate the posterior values of model parameters like tree topology and branch lengths conditional on the alignment data so as an output it provides a posterior probability distribution of the parameters of interest. This output of Bayesian analysis means built-in measures of uncertainties by providing posterior support for different tree topologies, branch lengths and other parameters. In this type of analysis the Bayes theorem takes the following form assuming a model M with parameters and an alignment X of N sites for S taxa: Pr(M|X) = Pr(X|M) x Pr(M)|Pr(X), where Pr(M|X) is the joint posterior probability distribution, Pr(X|M) is the likelihood function, Pr(M) is the prior probability and Pr(X) is the so called marginal likelihood or evidence.

If the model parameters are picked sensibly the expectation is that the Markov Chain of the Markov chain Monte Carlo (MCMC) algorithm will sample from the whole multidimensional posterior parameter space approximating the joint posterior distribution so the posterior probability of a parameter can be estimated with the frequency of sampled parameter values.

We have chosen the BEAST 2 package in order to show that Bayesian re-analysis of the data can provide added value to the biological interpretation even on the small set of SMPS somatic base substitution data^9^. In the BEAST 2 package there are a diverse set of substitution models and tree priors available that can be selected. Most importantly there are 4 different clocks models available in BEAST 2 including 3 different kind of relaxed and local clock methods besides the default strict clock.

## Results

### Strict clock versus relaxed clock

Branch lengths are calculated as the product of substitution rate variation and time elapsed in unconstrained analyses. Without fixing the rate parameter, branch length proportional to elapsed time cannot be estimated. As a terminological remark please note that in the context of the individual cell lineage tree we use substitution rate and mutation rate interchangeably. Traditionally the rate was assumed to be constant throughout the lineages giving rise to so called strict clock models^10^. But mutation rate can be different among lineages hence the so called relaxed clock models accommodated lineage specific rate variation and were applied in the context of traditional phylogenetic trees including several species. Many relaxed clock models have been implemented so far and different assumptions and parametric distributions have been used to draw samples for rates in the MCMC chain (for a good overview, see Heath and Moore 2014^11^). An important applied principle is to correlate a branch rate with its ancestral branch or keep them independent from each other and allow sudden shifts in rates in the tree.

BEAST 2 has four different clock models available out of the box currently, a strict clock, a relaxed uncorrelated exponential clock (UCED), a relaxed uncorrelated lognormal clock (UCLD) and a so called random local clock model (RLC).

BEAST 2 is able to co-estimate tree topology and lineage rate variation within the same runs so topology should not be necessarily fixed when estimating differences in branch specific mutation rates. We have applied the strict clock and the two uncorrelated relaxed clocks - UCED and UCLD - when sampling the joint posterior distributions of the Sanger Mouse Pilot Study. During unconstrained runs tree topologies – representing the consensus maximum clade credibility (MCC) topologies - remained constant, albeit with different level of posterior support, across the three clocks in both mice.

Both the unconstrained trees and the fixed MP trees were used for the MCMC runs estimating the mutation rate parameters and comparing different clock models to each other.

When investigating potential reasons to lean towards a strict or a relaxed clock interpretation of the SMPS data we have explored two ways:

1. UCLD model: coefficient of variation is indicative concerning how much variation among rates is implied by the data^12^ (see Drummond and Bouckaert, 2014, p147).
2. Model comparison/selection through path sampling^12^ (Drummond and Bouckaert, 2014, p139.).

#### UCLD

First, we looked at the marginal probability distribution of the coefficient of variation of the log-normal relaxed clock (UCLD). This coefficient is the standard deviation divided by the mean of the clock rate so basically the variance scaled by the mean rate. Strict clock can be ruled out if there is no appreciable amount probability mass near zero in the above-mentioned distribution. Figure 1 shows the marginal probability distribution of the coefficient of variation of the UCLD model for mouse_1 and mouse_2, respectively. The distributions were gained by combining together three runs with the same settings per mouse using the fixed MP tree. Almost identical distributions are sampled using unconstrained tree shapes (data not shown).

**Figure 1:**
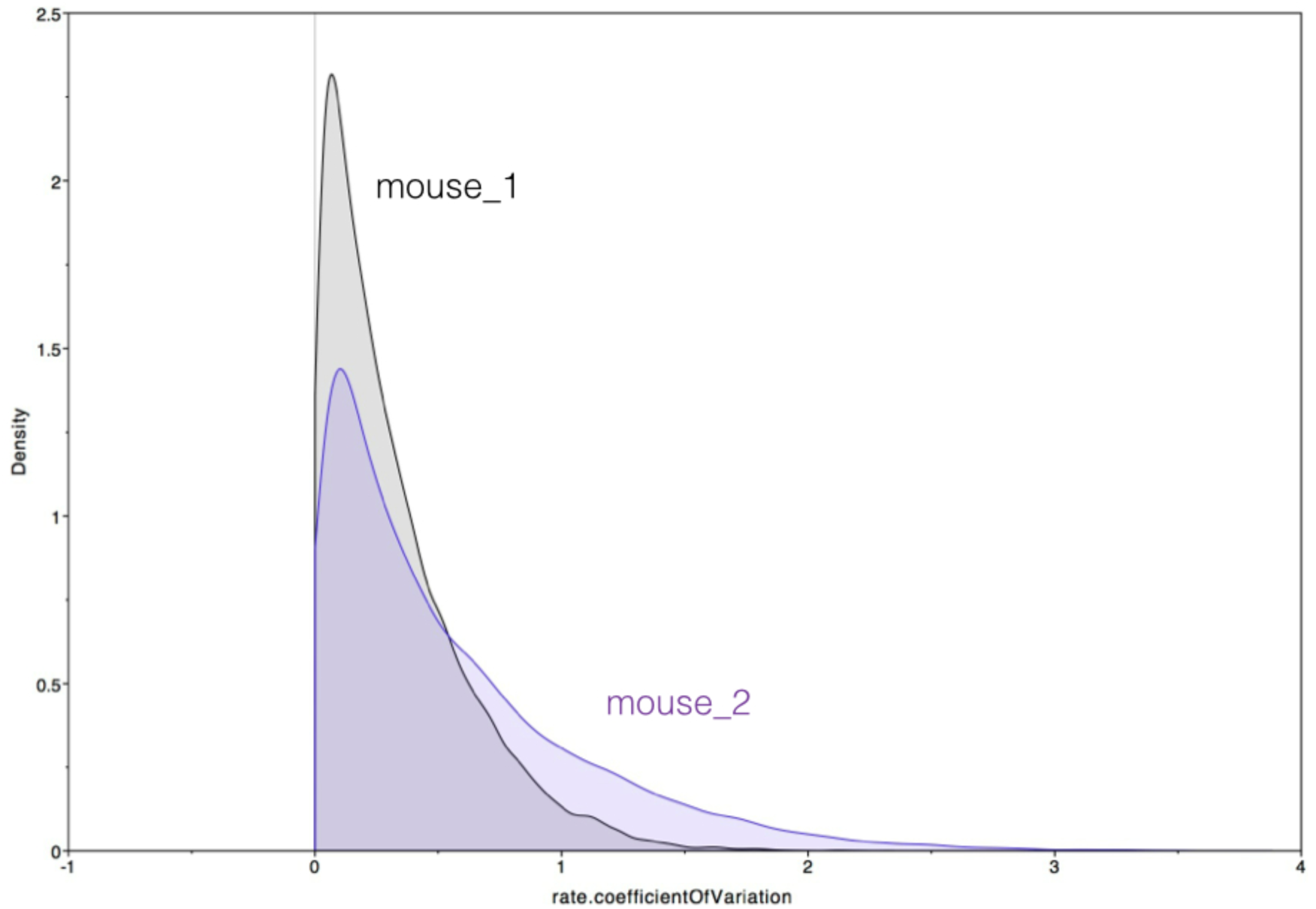
Marginal probability distribution of the coefficient of variation of the log-normal relaxed clock model in case of mouse_1 (black) and mouse_2 (blue) runs under fixed MP tree topologies, respectively.

Based on Figure 1, it’s obvious that in case of mouse_1 a big chunk of the probability mass of the coefficient of variation is around zero so the data cannot be used to reject a strict clock. For mouse_2 there’s still a significant amount of the probability mass close to zero although there’s more mass falling between 0.5 and 2 than for mouse_1. To put it another way, the posterior mean coefficient of variation is 0.318 for mouse_1 and 0.546 for mouse_2 which means that the rate is varying by 32 (mouse_1) or 54% (mouse_2) in different clades over the tree. The >50% rate variation suggests that indeed at least for mouse_2 the data allows to consider actually different mutation rates along some lineages.

Next we looked at the mean rates and 95% highest posterior density (HPD) intervals UCLD is producing. The interval is the so called highest posterior density interval that includes the smallest, most compact range of values, in this case rate values, amounting to the n %, in this case 95 %, of the posterior probability mass. It is considered by many the Bayesian analog of the confidence interval. As such, the 95% HPD interval can indicate how much uncertainty there is in these assigned rates. Figure 2 and Figure 3 show the mean rates and 95% HPD intervals under the UCLD model for the two mice, respectively, assuming a fixed MP tree topology.

**Figure 2:**
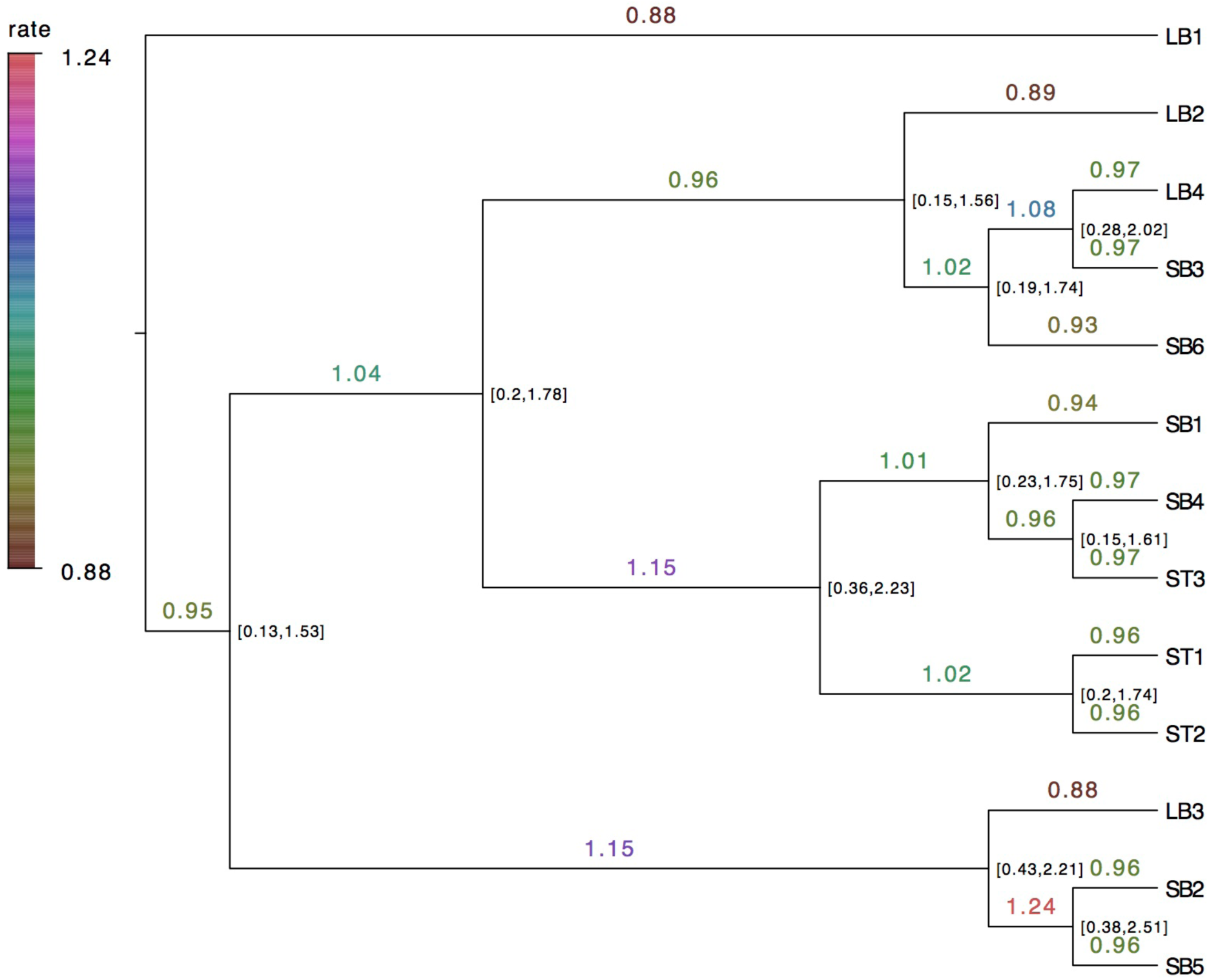
Mean sampled mutation rates using UCLD clock model assuming a fixed MP tree for mouse_1. Branch labels show mean mutation rates and node labels provide the 95% HPD interval (see text). Values are rounded to 2 decimal places. The colored bar shows the color codes of the mean rates.

**Figure 3:**
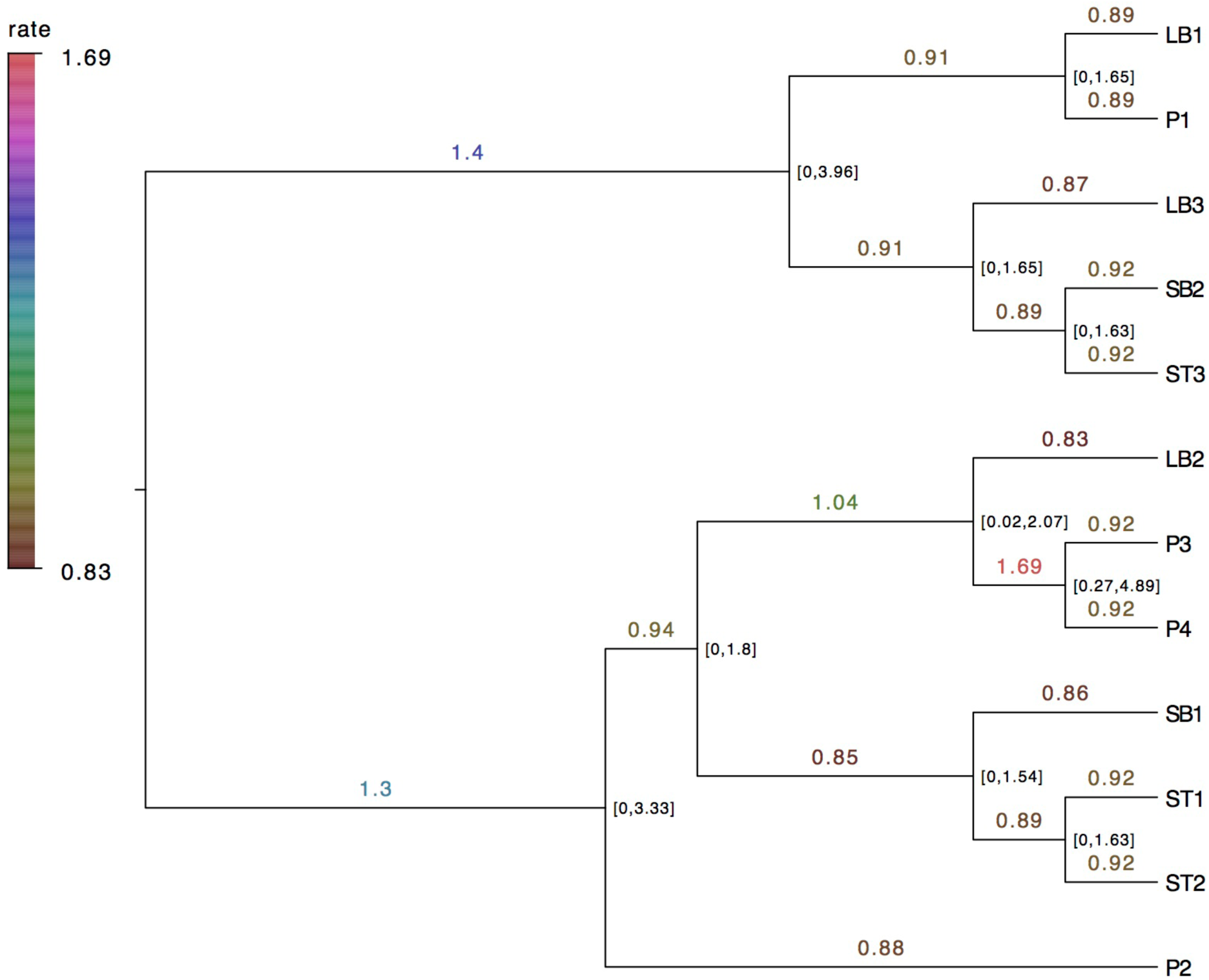
Mean sampled mutation rates using UCLD clock model assuming a fixed MP tree for mouse_2. Branch labels show mean mutation rates and node labels provide the 95% HPD interval (see text). Values are rounded to 2 decimal places. The colored bar shows the color codes of the mean rates.

Looking at the 95% HPD ranges it is apparent that there’s a lot of uncertainty in the sampled rates for both mice. Since the numbers were rounded up to two decimal places the smallest values, like 0.001 or 0.003, are displayed as 0 actually. The rate values are provided as units of substitutions per site per unit time since no absolute divergence time estimation have been performed. The broad HPD ranges suggest that one need to be careful with any possible interpretations concerning the rates. Concerning mouse_1, the mean rates range between 0.88 - 1.24 so within 25% of 1. The only outlier mean rate, 1.24, belongs to the branch uniting the identical SB2 and SB5 sequences, SB2 representing a proximal small bowel crypt and SB5 representing a distal one. Since the coefficient of variation supported the strict clock, this outlier is more like a curiosity than a signal in the UCLD context. Mouse_2 though presents one spectacular outlier branch rate of 1.69 forming a clade leading to the identical P3 and P4 organoids representing single cells from the dorsolateral and ventral part of the prostate, respectively. The 1.69 mean rate is twice as much as the lowest mean rates calculated and this reflects the posterior mean coefficient of variation value of 0.546 calculated for mouse_2 suggesting a rate varying by 54% in different clades over the tree. Since the value of the coefficient gives enough reason to lean towards the UCLD relaxed clock model this creates a more confident finding in mouse_2. Supplementary Material section entitled ‘*Mean sampled mutation rates using the UCED clock’* contains the trees and sample mean mutation rates under the UCED relaxed clock model.

#### Model selection via Bayes Factor

Table 1 shows log marginal likelihood (ML) estimations by path sampling for the three clock models used in unconstrained settings or with a fixed MP tree for the two mice with other parameters remaining the same. ML estimates using 32 steps turned out to be no different than those with 16 steps.

**Table 1:** Log marginal likelihood estimates using path sampling for fixed MP trees and unconstrained trees for mouse_1 and mouse_2. The values displayed there are the mean values of three technical replicate path sampling runs within the same setting/row. Besides the clock models, other models and parameters were set constant: the reversible-jump based model was used for substitution model, coalescent tree prior with constant population was used for the branching process. The marginal likelihood values can be used to assess the fit of different models to the same data via Bayes Factor calculation. The corresponding values highlighted in bold, italicized text provide support to one model over the over.

|  | Strict,<br>32 steps | Strict,<br>16 steps | UCED,<br>32 steps | UCED,<br>16 steps | UCLD,<br>32 steps | UCLD,<br>16 steps |
| --- | --- | --- | --- | --- | --- | --- |
| Mouse_1<br>Fixed MP tree | -119.5 | -119.61 | -119.5 | -119.49 | -119.4 | -119.4 |
| Mouse_1<br>Unconstrained | -141.3 | -141.14 | -140.9 | -141.24 | -141.2 | -140.69 |
| Mouse_2<br>Fixed MP tree | **-77.1** | **-77.1** | **-75.9** | **-76.0** | -76.7 | -76.66 |
| Mouse_2<br>Unconstrained | **-92.6** | **-92.68** | **-91.4** | **-91.16** | -92.3 | -92.17 |

In BEAST 2 the reported marginal likelihoods (ML) are in the log-space and consequently for Bayes Factor calculation the respective ML-s of competing models can be subtracted from each other. The corresponding values on Table 1 highlighted in bold, italicized text provide support to one model over the over. Taking only the 32 step results for mouse_2, the fixed MP tree setting gives lnBF = -75.9 - (- 77.1) = 1.2 and that is in the 1.1 -3 range providing positive, albeit not particularly strong, support for the UCED relaxed clock over a strict clock assumption^12^. The biggest difference between the 3 technical replicates within the same setting were all much smaller than the difference between the means of the two clock settings compared to each other - 0.2 vs 1.2 for mouse_2 unconstrained and 32 steps, 0.05 vs. 1.2 for mouse_2 fixed MP tree and 32 steps - making it comfortable to calculate the Bayes Factor.

On the other hand for both mouse_1 settings, the differences between the mean marginal likelihood values of the different clock setting replicates are negligible and all less than 0.5 so one model cannot be comfortably selected over the other using Bayesian tools.

Substitution rates can be different across different sites within the same sequence and the gamma model has been widely used to fit the data and it is also available in BEAST 2^13^. The models without gamma rate heterogeneity have consistently outperformed the corresponding models with gamma rate heterogeneity added, please see Supplementary Material, section entitled ‘*Gamma rate heterogeneity*’.

## Discussion

The Bayesian re-analysis of the SMPS cell lineage tree data with the BEAST 2 package allowed us to extend the interpretation space of the underlying mechanisms of cell lineage tree formations. On one hand, BEAST 2 provided support for topologies similar to the original Maximum Parsimony analysis, the differences favoring the perfect phylogeny achieved with Maximum Parsimony. Most importantly, unbiased clock analysis with BEAST 2, unavailable for other tree reconstruction methods, made it possible to investigate clock-like behaviour and provide, for the first time, different mutation rate estimations of different intraindividual cell populations and lineages. Table 2 provides the following pro and contra arguments concerning the choice between strict or non-strict clock models for the data.

**Table 2:** Two ways to argue for the clocklikeness for the SMPS data. Path sampling is the standard way to do model comparison and selection based on marginal likelihood and Bayes Factor calculation. The distribution of UCLD’s coefficient of variation is another way to argue for or against strict clock depending the amount probability mass near zero.

|  |  |  |
| --- | --- | --- |
| Method used | mouse_1<br>Marginal likelihood | mouse_2<br>Marginal likelihood |
| Path sampling | Undecidable -> Strict | Relaxed, UCED |
| UCLD:<br>coefficient of variant | Strict | Relaxed |

Although path sampling could not discriminate between clock models in case of mouse_1, since the strict clock model is using less parameters and is more simple than relaxed clocks, favouring the strict clock is the economical – but not necessarily correct - choice over relaxed clocks.

While for mouse_1 strict clock is the preferred option, for mouse_2, rate variation - at least along one branch of the tree, but potentially more - might be reasonably and preferably assumed based on this type of BEAST 2 clock analysis.

The near equal performance of strict clock and relaxed clock models is ambiguous, informative and promising at the same time. In mouse_2 the value of the coefficient of variation gives enough reason to lean towards the UCLD relaxed clock model and path sampling also supports the UCED model over the strict clock.

Interestingly, a clade of two prostate organoid lines in mouse_2 presented one spectacular outlier branch rate compared to the mean rates calculated. This is promising as in mouse_2 the possibility of rate variation along a branch opens up the space for subsequent clock analysis on extended single cell resolution lineage data to motivate a biological interpretation for the possible elevated mutation rates in prostate or other lineages. For instance a recent study on three prostate cancer patients found that somatic mutations were present at high levels in morphologically normal, “healthy” prostate tissue as well, distant from the cancer site^14^. This is another hint that the prostate is a hotbed of unusual/abnormal mutational processes. Perhaps a higher default tissue specific mutational rate in the prostate might be part of the explanation of high prostate cancer incidence amongst different type of cancers?

As we mentioned earlier, the rate values are provided as units of substitutions per site per unit time (SST) since no absolute divergence time estimation has been performed. One question that might be asked is whether this rate estimation can be expressed per cell division as units of substitutions per site per cell division. The sampled endodermal tissues are all mitotically active tissues so it is reasonable to assume that DNA replication tied to subsequent cell divisions is a major cause of the observed somatic base substitutions. While the thorough examination of cell division normalised mutation rates is out of the scope of this paper the calculation could go like this: if there are on average X cell divisions since the origin of the tree in a particular tissue, then going from Y substitutions per site per unit of time (SST) to Z substitutions per site per cell division (SSD), just use Z = Y / X. However, as far as we know, there are no strict estimations in mice on the tissue specific number of cell divisions happening throughout the life of a mouse. One exception is small bowel crypt cells where for instance small-bowel Lgr5-positive stem cells have been reported to divide every 21.5 hours^15^. So for instance for mouse_1 (died 116 weeks old) there are two identical tips SB2 and SB5 with four, subsequent branches leading to them from root with four different mean sampled mutation rates (calculated from the root, see Supplementary Fig. S7): 0.81 -> 1.66 -> 1.98 -> 0.83. Based on this SSD can be calculated for the last branch leading to the two tips SB2 and SB5 with 0.96 mean sampled rate by using age and division rate of SB cells which is 21.5 hours: (116 (weeks) * 7 (days) * 24 (hours)) /21.5 (hours/division) = 906.4186 ~ 900 divisions throughout 116 weeks and this number can be used to calculate with the help of 0.96 SST as 0.96 SST / 900 divisions ~= 0.001 substitutions per site per cell division (SSD). 900 cell divisions in the same stem cell niche sounds incredibly big. Nevertheless 900 divisions in total might still be a an acceptable ballpark estimation for adult mouse small intestine epithelial cells if we consider that the turnover time for one layer of epithelial cells is about 3-5 days, meaning that those epithelial cells of the intestinal villi are completely replaced within that time. Maintaining this food and nutrient absorbing function at the same level is essential for the body so the stem cell niche behind the epithelial cells is going through cell divisions on a daily basis for 2 years.

Relaxed clock models differ in the underlying process of rate variation assumed or aimed at capturing; it can be thought of as a gradual process that is captured with autocorrelated rate models or episodic, sudden rate changes as in case of the independent rate models, out of which we have tried UCED and UCLD. Rate autocorrelation in case of the usually investigated inter-species trees can happen due to genetic factors determining rate variation. Concerning organismal cell lineage trees there might be a reason to develop a new variant of the relaxed clock model. In the clinical setting lineage rate variation might be applied to single cell resolution tumour and cancer samples exhibiting much higher default mutation rates than healthy tissues.

Based on this study we have reason to assume that unbiased clock analysis applied to single cell resolution cell lineage tree data adds a new, useful layer to our understanding of cell lineage trees. This understanding can also lead to the discovery of new biological principles at work during the normal life cycle of multicellular organisms and might also provide actionable insights in medicine.

## Methods

### BEAST workflow

Figure 4 below shows the methodological workflow followed throughout the analysis.

**Figure 4:**
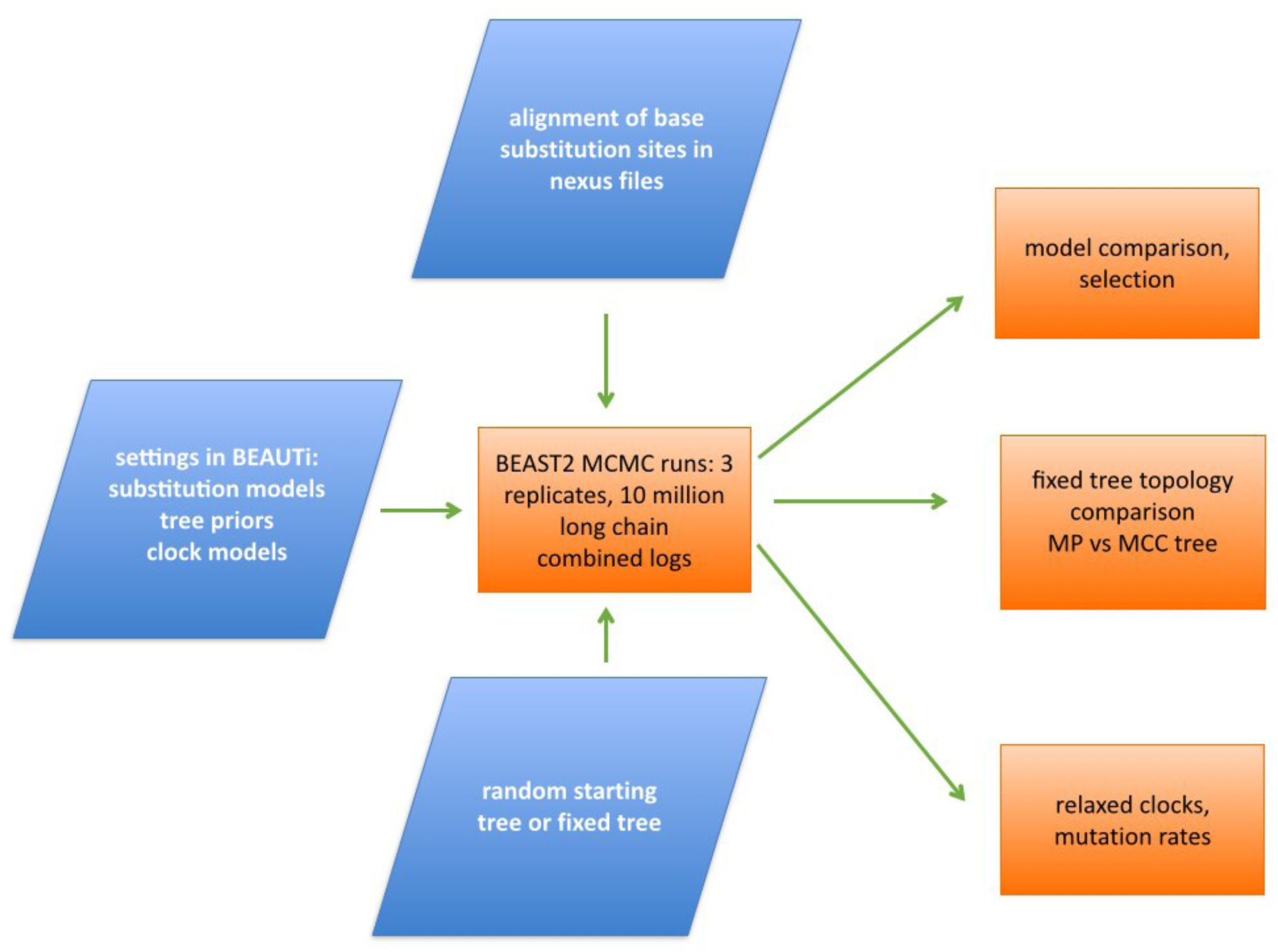
Methodology workflow. BEAST 2 components applied throughout the analysis from importing alignment, setting up model parameters, MCMC runs to rate analysis, topology comparison and model selection.

Supplementary Table 2 of the SMPS study containing the 35 embryonic mutations for the two mice have been used to assemble the alignments of sequences in nexus files for the two mice separately^6^. BEAUTi 2, available from the BEAST 2 software package, was used to import the alignments and set up the models, priors and parameters like substitution site models, tree priors, clock models amongst others. The output of BEAUTi is a BEAST xml file that is the input of the BEAST runs. In case of the fixed tree topology sampling experiments, like MP trees, the BEAST xml files have been manually changed to provide the fixed trees in newick format and the four operators changing the tree topology - SubtreeSlide, narrow, wide, WilsonBalding - have been turned off. BEAST 2 MCMC runs have been set up as follows: the default 10 million length Markov chains proved to be robust enough to provide a sufficient effective sample size (ESS) of independent samples in the range of several hundreds but usually thousands individually and sufficient enough for the parameters estimated. A burn-in of 10% has been cut off before analysing the results further. Three technical replicates have been produced with every chosen settings starting from different random seeds and then posterior values have been compared alongside the technical replicates in order to check for convergence. For calculating relaxed clock mutation rates tree-logs from all three replicates have been merged. LogCombiner, available within the BEAST 2 software, was used to merge trace logs and tree-logs and apply 10% burnin cutoff. FigTree v1.4.1 (http://tree.bio.ed.ac.uk/software/figtree/) and DensiTree v2.0 have been used to visualise annotated trees^16^. BEAST 2 version 2.1.3 has been used throughout the study.

### Model comparison and selection

In BEAST 2 there are many model parameters that can be selected from substitution models to branching process/tree priors, population parameters and clock models. Since our focus was on testing different clock models first we needed to do unconstrained MCMC runs without a fixed tree topology and then to apply model comparison and selection methods to decide which parameters will be used. As Baele and Lemey put it^17^ referring to Steel^18^: *“The aim of model selection is not necessarily to find the true model that generated the data, but to select a model that best balances simplicity with flexibility and biological realism in capturing the key features of the data”* In the lack of custom tailored models adjusted to the specific features of organismal cell lineage trees we have mainly relied on what’s available in BEAST 2 and what can be picked based on model selection. As far as substitution models are concerned we have used a reversible-jump based model, see section entitled ‘*Substitution model selection*’ below. For the branching process prior model selection settled down on using the coalescent tree prior with constant population, see ‘*Tree prior selection*’ section below. Bayes factor evaluation is the standard way to perform model selection in Bayesian phylogenetics, the Bayes Factor being the ratio of two marginal likelihoods

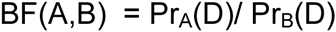

for the two models, A and B, under comparison. BF > 1 supports model A over B. Calculating the marginal likelihood is challenging as it involves integrating over all possible model parameters. Many methods have been developed out of which we have used path sampling, briefly discussed in section entitled ‘*Bayes factor calculation of clock models via path sampling*’ below.

### Substitution model selection

DNA substitution models describe nucleotide substitutions over time by a continuous-time Markov model with instantaneous rate matrices and 12 possible transition rates. There are many models available starting from the simplest one (Jukes and Cantor) to more complex ones. The complex models use different weighted parameters for transitions (A <-> G, C <-> T) and transversions (purines <-> pyrimidines) and allow different base frequencies amongst others.

Instead of using Bayes Factor based model comparison and selection with the numerous substitution model variants available in BEAST 2 we have used the reversible-jump based model (RB) add-on from the RBS package ^19^. Please see Supplementary Material section entitled *Reversible-jump based model for substitution model selection* for details.

### Tree prior selection

The main question here was to decide between the default Yule tree prior, a pure birth process, or some other variants of it, a Birth Death model for instance, and the coalescent tree prior. The coalescent process comes from population genetics and deals with gene trees tracing back the allele variants into their most recent common ancestors. Since the model is using population parameters by default we have used the coalescent tree prior with a constant population. See Supplementary Material section entitled *Tree Prior Selection: coalescent models and Skyline plots* for details.

### Bayes factor calculation of clock models via path sampling

Path sampling is a computationally intensive method consistently outperforming importance sampling methods like the HME or the AICM approach^20^. In path sampling the marginal likelihood is estimated by sampling from the product of the prior and the likelihood under a hyperparameter due to which the MCMC chain samples from between the prior and the posterior.

In order to sample from sensible values and avoid the parameters to escape to infinity the population size prior has been adjusted. Instead of using the improper 1/X that might lead population size to zero a lower and upper bound was set on population size, denoted here as [m/10, 10*m] where m is the mean population size sampled from the posterior distribution after the initial MCMC runs with the corresponding settings. Log marginal likelihood estimations were calculated as the mean values of three technical replicate path sampling runs within the same setting. By default 32 steps were chosen and the MCMC chain length was 300000. In order to get reassurances that the path sampling analysis is actually correct, the process was repeated with 16 steps so ML estimates can be compared together.

## Acknowledgements

The authors want to acknowledge Botond Sipos for suggesting the idea of applying relaxed clock models to cell lineage data. The authors would also like to thank Jakub Truszkowski, Nick Goldman and Henning Hermjakob for the critical feedback provided.

## Author Contributions Statement

A.Cs. and R.B. designed the simulations. A.Cs. performed the simulations. A.Cs. and R.B. analysed the data. A.Cs. wrote the manuscript. Both authors reviewed and approved the manuscript.

## Competing Interests

The authors declare no competing financial interests.

## Supplementary material

### Topology

Unconstrained BEAST 2 MCMC runs with random tree sampling have been performed in order to decide how the maximum parsimony tree topology is supported. The reversible jump substitution model (Bouckaert et al. 2012) was used as substitution model, without gamma rate heterogeneity, and the coalescent tree prior was used with constant population as a tree prior, see details in the *Methods* section.

There are many flavours of summary or consensus trees of sampled posterior trees out of which we are using the so called maximum clade credibility tree (MCC from now on) which is the default setting in TreeAnnotator or DensiTree and was shown to perform well on a range of criteria^1^. The MCC tree is the sampled tree with the maximum product of posterior clade probabilities.

Supplementary Figure S1 and Supplementary Figure S2 shows the posterior support and the topology differences in case of mouse_1 and mouse_2, respectively.

**Supplementary Figure S1:**
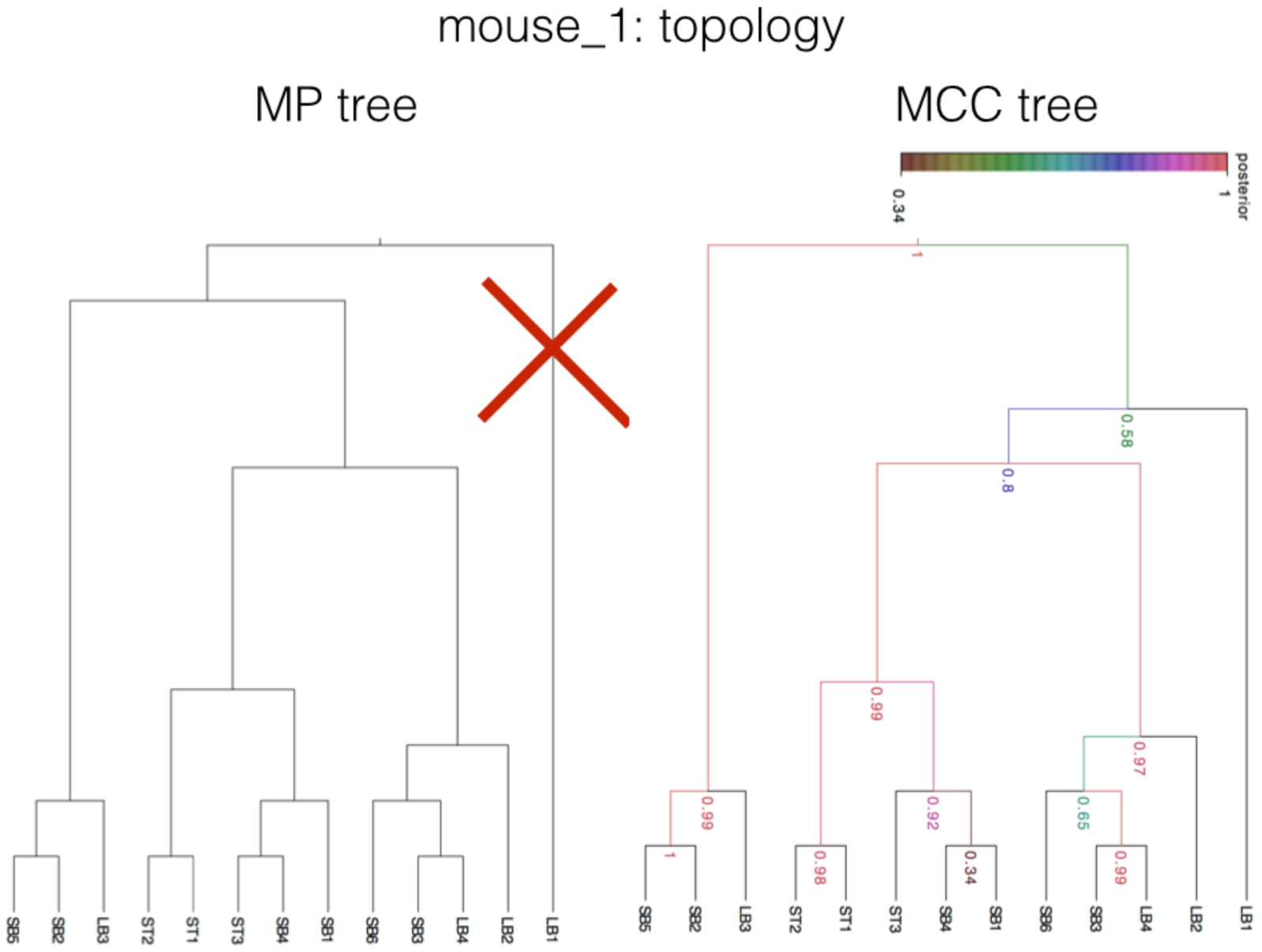
Mouse_1, comparing the maximum parsimony (MP) tree with the maximum clade credibility (MCC) tree. The cross in the MP tree highlights the difference in topology. The numbers on the nodes on the MCC tree denote posterior support for the particular clade,the coloring highlights the amount of posterior support for particular clades. Percentage support can be gained by multiplying the numbers with 100.

**Supplementary Figure S2:**
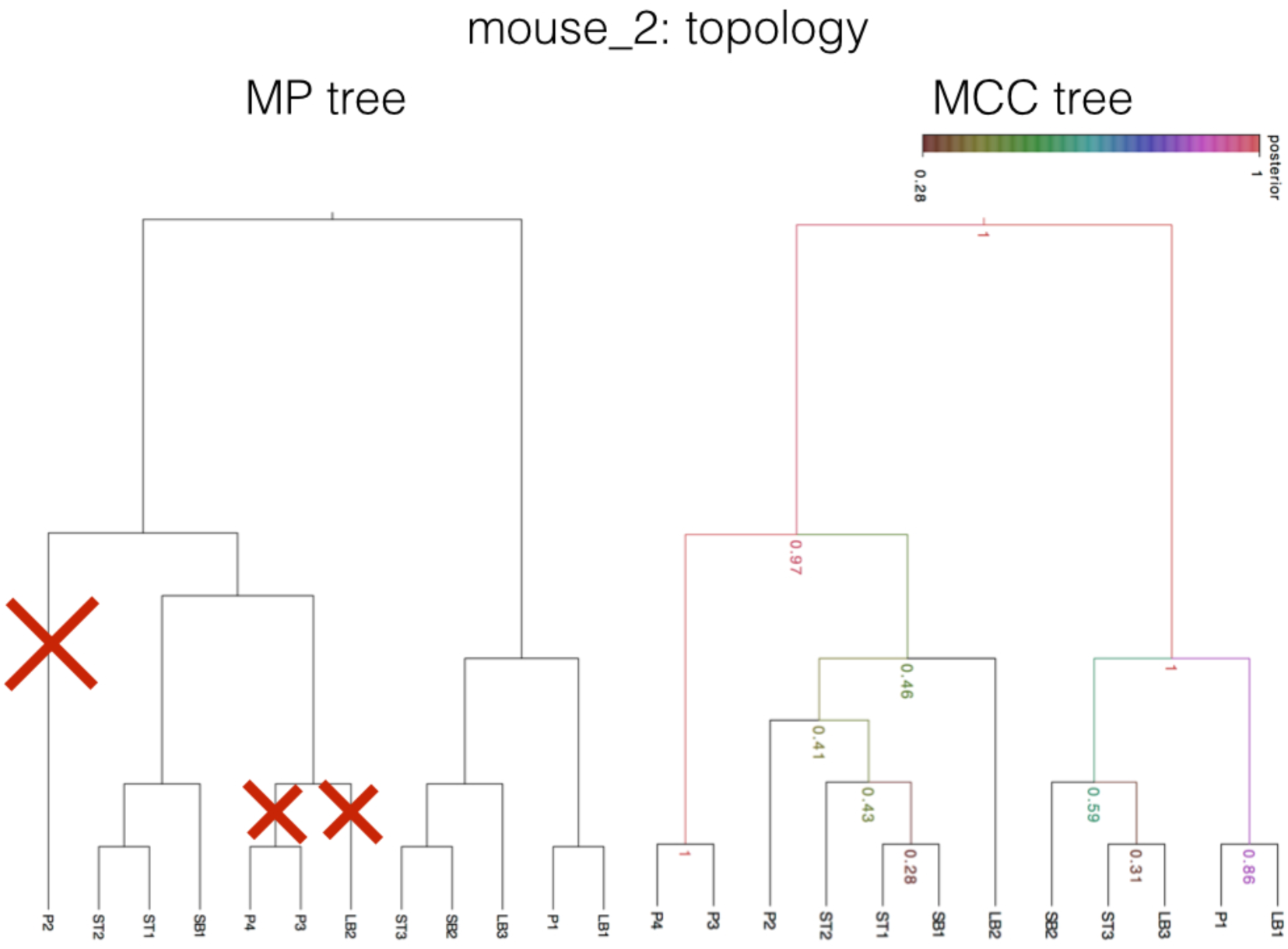
Mouse_2, comparing the maximum parsimony (MP) tree with the maximum clade credibility (MCC) tree. Crosses in the MP tree highlights the differences in topology. The number on the nodes on the MCC tree denote posterior support for the particular clade, the coloring highlights the amount of posterior support for particular clades. Percentage support can be gained by multiplying the numbers with 100.

Concerning mouse_1 more than 50% posterior support is provided for the majority of the clades in the maximum parsimony reconstruction. Not all clades have this majority support, for instance, clade (SB1,SB4) containing identical sequences only has 34% support. In mouse_2 the posterior support is more than 40% for the majority of the clades excluding clades containing leaves with identical sequences. These latter clades include (SB1,ST1) with 28% or the (LB3,ST3) clade with 31% support. Topology-wise the only remarkable difference in mouse_1 is the placing of the ancestry of LB1. In mouse_2 there are major differences in 3 clades containing 4 organoids, P2 closest to the root, LB2 and the identical P3 and P4.

The question is how to explain or reconcile cell lineage tree shape in clades differently reconstructed by the MP and Bayesian approaches. Concerning the MP reconstruction (Behjati et al. 2014) *“A unique most-parsimonious solution was found for each mouse into which all embryonic variants fitted with no homoplasy.”*

Homoplasy can mean two different phenomena in this context: i., **convergent evolution** when the same somatic base substitutions appear in tips belonging to two different clades and not present in their last common ancestor and ii., **back mutations** or reversals when there is a sequence of X -> Y -> X back and forth base substitutions along a clade in the tree, for instance an C -> T -> C pattern occurring as a result of subsequent divisions.

Supplementary Figure S3 below shows how the MCC tree topology can explain the placement of LB1 in a different clade with the help of the two types of homoplasies mentioned before. Scenario A contains 2 homoplasies, 1 convergence (2 convergent events) and a back mutation while Scenario B includes only convergence with 2 converging base substitution events.

**Supplementary Figure S3:**
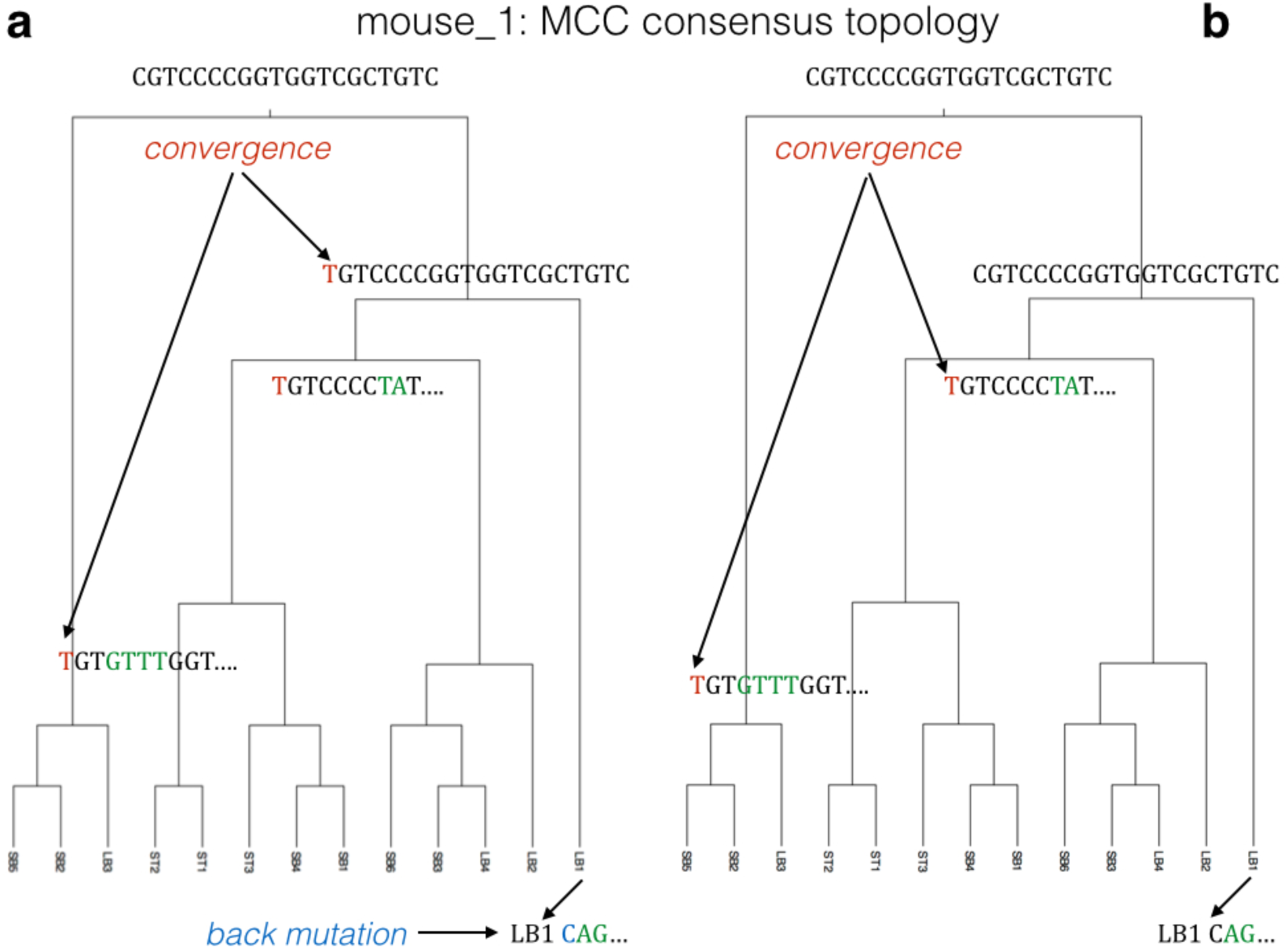
Unlikely hypothetical homoplasy scenarios backing the maximum clade credibility (MCC) consensus tree topology for mouse_1. **(a)** Scenario A contains the co-occurrence of 2 convergent mutation events and a back mutation. (**b)** Scenario B includes only convergence with 2 converging base substitution events. Color code: red: convergent base substitution, blue: back mutation/substitution, green: somatic base substitution without homoplasy.

Supplementary Figure S4 below shows how the consensus MCC tree topology can be explained with homoplasies in case of mouse_2. Scenario A involves a back mutation and a convergence while Scenario B includes 2 convergences out of which one is the hypothetical co-occurrence of 3 converging base substitution events along a cell lineage tree.

**Supplementary Figure S4:**
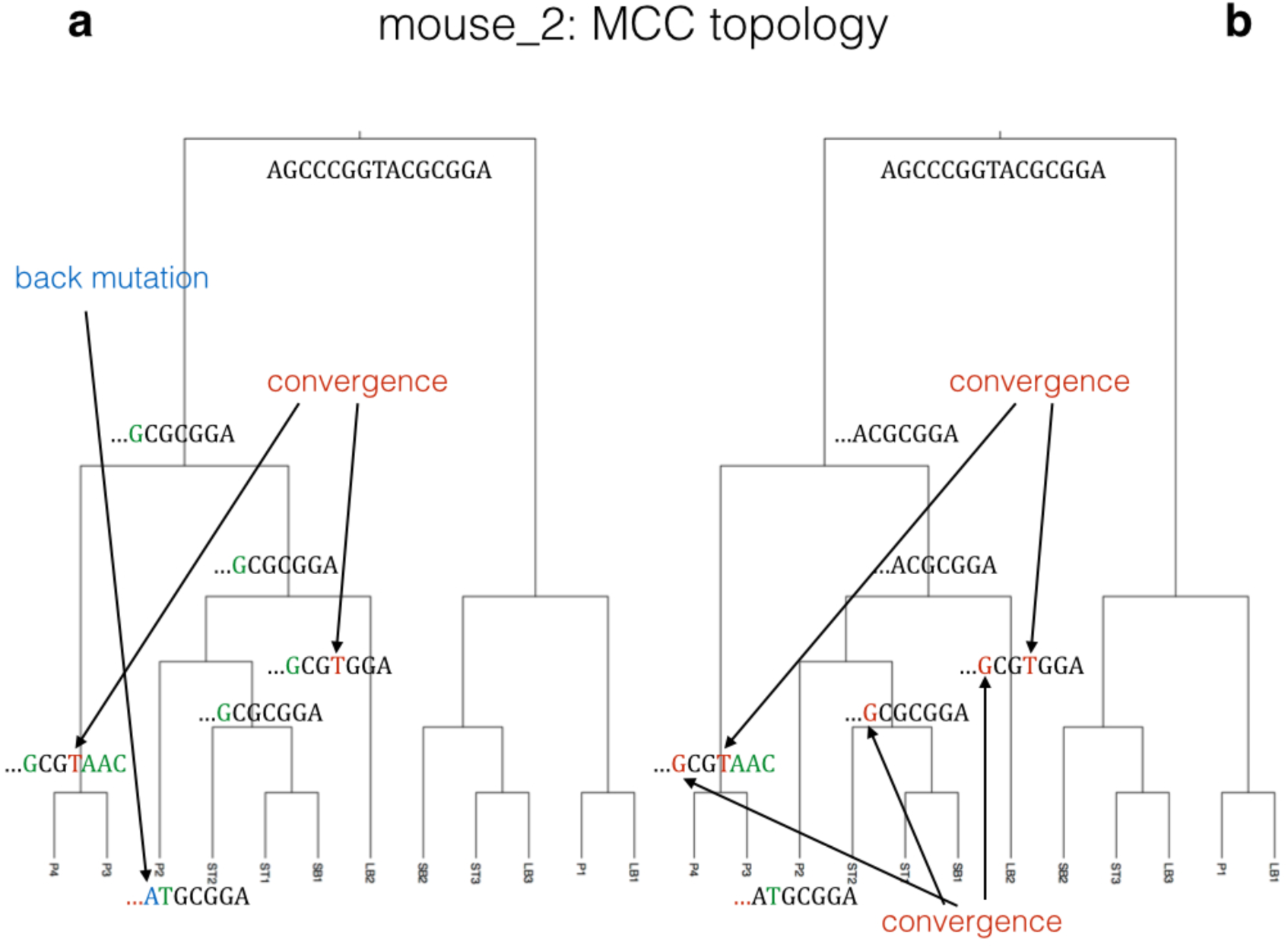
Unlikely hypothetical homoplasy scenarios backing the maximum clade credibility (MCC) consensus tree topology for mouse_2. **(a)** Scenario A contains the co-occurrence of 2 convergent mutation events and a back mutation. (**b)** Scenario B includes only convergence with 2 converging base substitution events. Color code: red: convergent base substitution, blue: back mutation/substitution, green: default somatic base substitution.

The probability of any kind of homoplasy occurring within a mouse cell lineage tree is considered to be extremely small. If the probability of 1 mutation occurring is already small, say 10^−9^/base*year^2^ then the probability of a back mutation happening in a subsequent cell division will be even less and the probability of no substitution happening will be even closer to 1. The same applies in case of convergence events when 2 mutations occur at the same site irrespectively of the order of these mutations happening. When the data admits a perfect phylogeny without homoplasies of any sort it is unlikely that the perfect phylogeny is wrong, or suboptimal, under any reasonable substitution model of evolution. Nevertheless the sites were particularly selected for variability in the SMPS.

Note that for mouse_1 the Bayesian reconstruction supports the MP topology with the exception of LB1. For mouse_2 the Bayesian reconstruction provides >40% for most of the clades and ~30% support for 2 clades closer to the leaves.

In order to show that there is enough data to establish the tree topology we have sampled from the prior distributions only with BEAST 2 and showed that without using the sequence data the shapes of the trees for mouse_1 and mouse_2 collapse with less than 6.5% posterior support for the particular clades. See next section in Supplementary Material, entitled ‘*Sampling from the prior*’.

### Sampling from the prior

In order to show that there is enough data to establish the tree topology we have sampled from the prior distributions only with BEAST 2 and showed that without using the sequence data the shapes of the trees for mouse_1 and mouse_2 are collapsing with less than 6.5% posterior support for the particular clades. As opposed to this when sequence data was used during posterior sampling it firmly suggested preferable clade topologies in case of mouse_1 This can be illustrated by showing the MCC tree showing the tree sampled with the maximum product of the posterior clade probabilities, thereby summarising the posterior support on different clades. This support is more than 50% for all of the clades with leaves containing different sequences. Please see Supplementary Figure S5 comparing posterior tree topology support with and without sequence data for mouse_1. Mouse_2 MCMC runs with sequence data provided suggested preferable clade topologies. This support was more than 40% for all of the clades excluding clades containing leaves with identical sequences. Please see Supplementary Figure S6 comparing posterior tree topology support with and without sequence data for mouse_2.

**Supplementary Figure S5:**
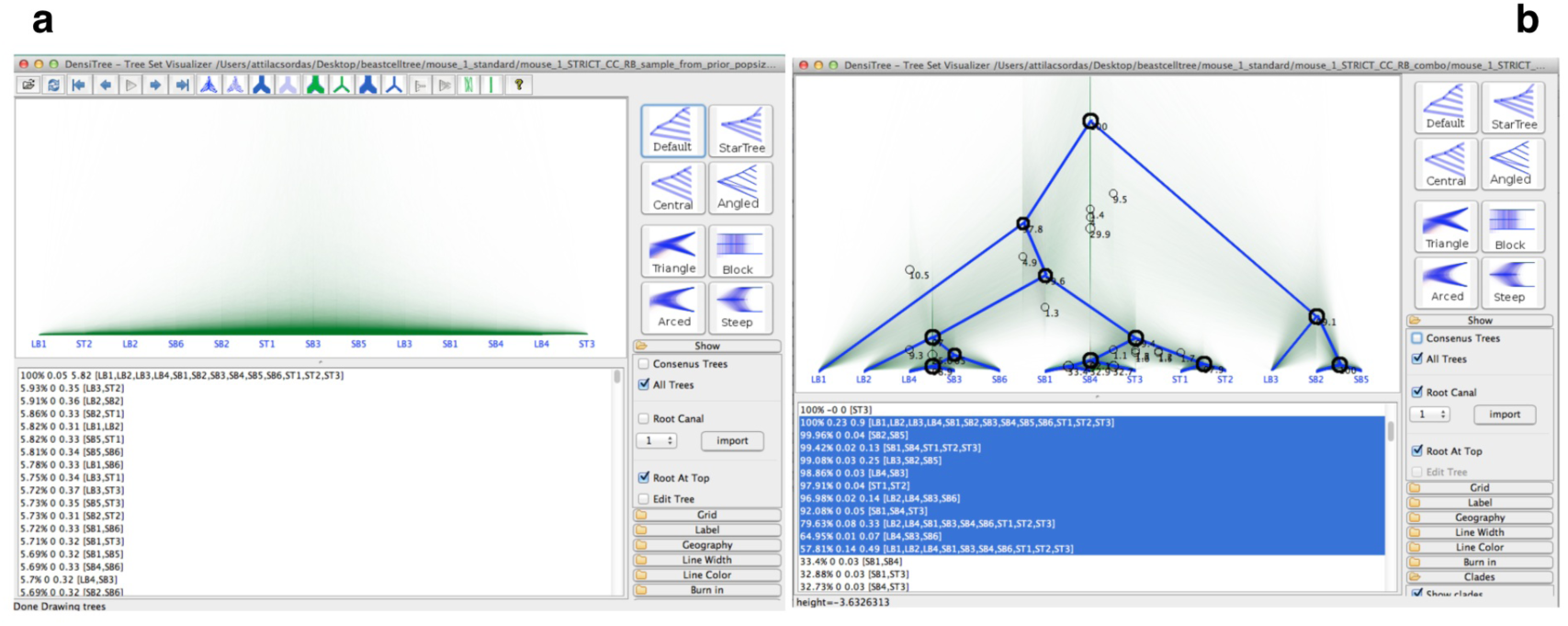
Sampling from the prior distribution only versus sampling from the priors + data in case of mouse_1. **(a)** Sampling from the prior distribution only is not sufficient to establish a tree topology in case of mouse_1 as all of the sub-clades have been sampled less than 6% of the time individually. **(b)** When sequence data was used during posterior sampling it firmly suggested preferable clade topologies. This can be illustrated by showing the MCC tree showing the tree sampled with the maximum product of the posterior clade probabilities, thereby summarising the posterior support on different clades. This support is more than 50% for all of the clades with leaves containing different sequences.

**Supplementary Figure S6:**
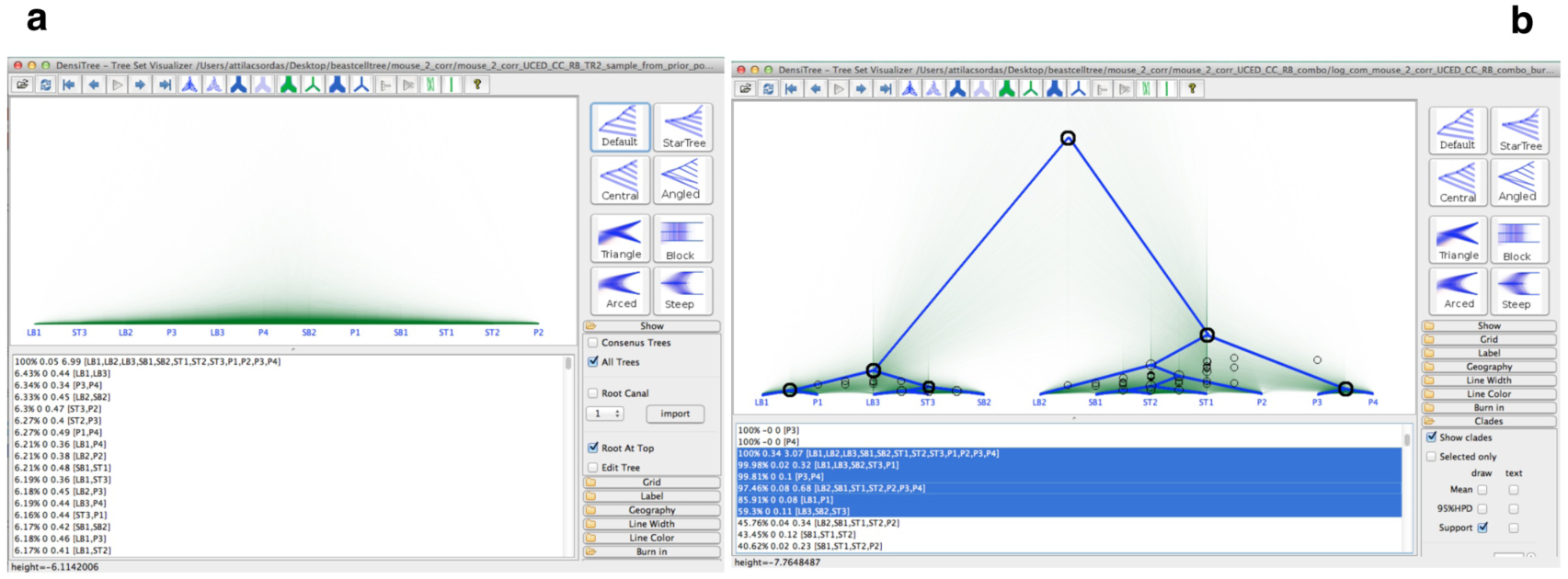
Sampling from the prior distribution only versus sampling from the priors + data in case of mouse_2. **(a)** Sampling from the prior distribution only is not sufficient to establish a tree topology in case of mouse_2 as all of the sub-clades have been sampled less than 6.5% of the time individually. **(b)** When sequence data was used during posterior sampling it suggested preferable clade topologies, posterior support for the clades summarised by the MCC tree. This support was more than 40% for all of the clades excluding clades containing leaves with identical sequences.

### Mean sampled mutation rates using the UCED clock

Supplementary Figure S7 and Supplementary Figure S8 shows the mean sampled rates and 95% rate highest posterior density intervals assuming the fixed MP tree under the UCED model for mouse_1 and mouse_2, respectively. Supplementary Figure S9 shows the same for the unconstrained mouse_2 run sampling rates under the UCED clock.

**Supplementary Figure S7:**
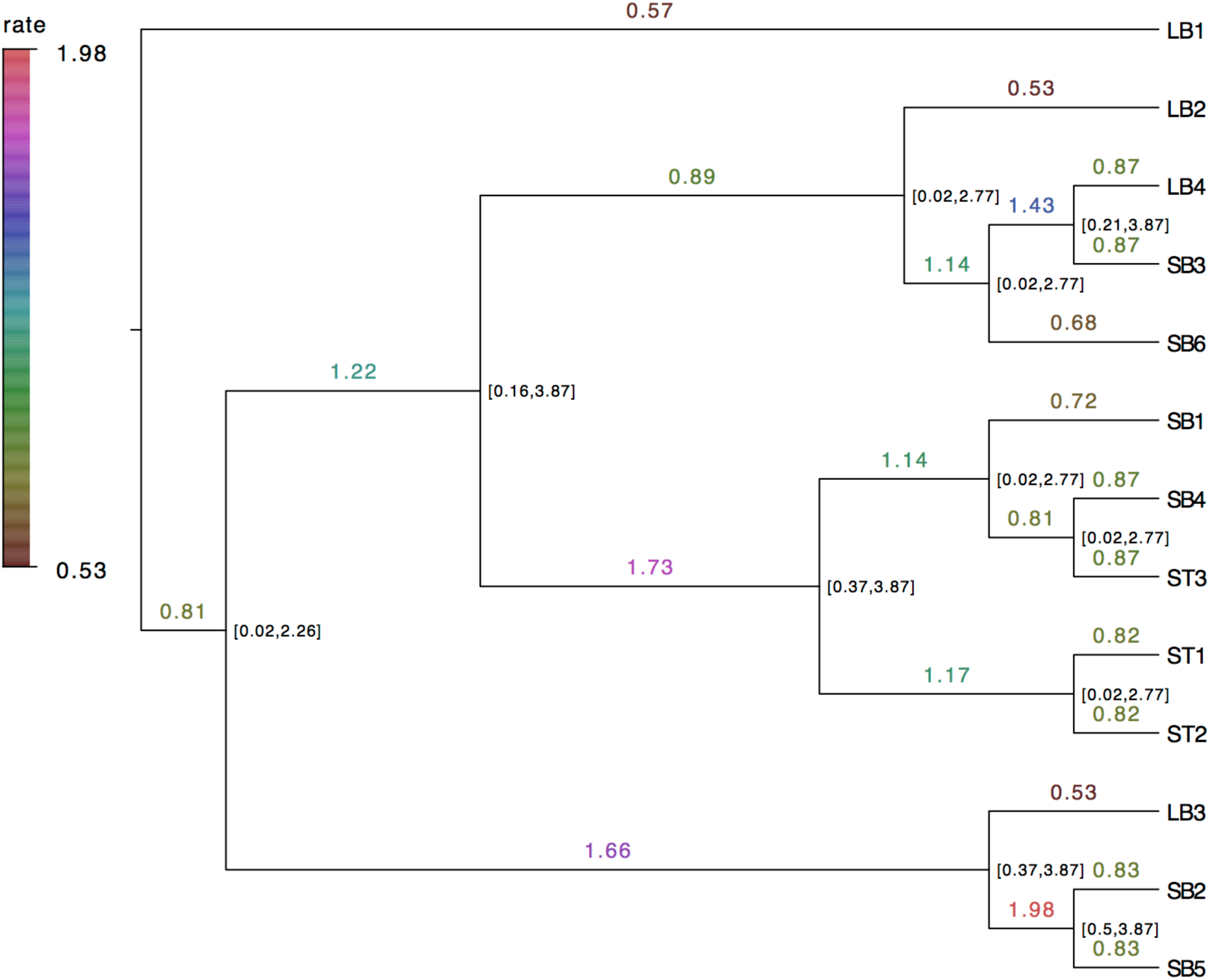
Mean sampled mutation rates using UCED clock model assuming a fixed MP tree for mouse_1. Branch labels show mean mutation rates and node labels provide the 95% HPD interval (see text). Values are rounded to 2 decimal places. The colored bar shows the color codes of the mean rates.

**Supplementary Figure S8:**
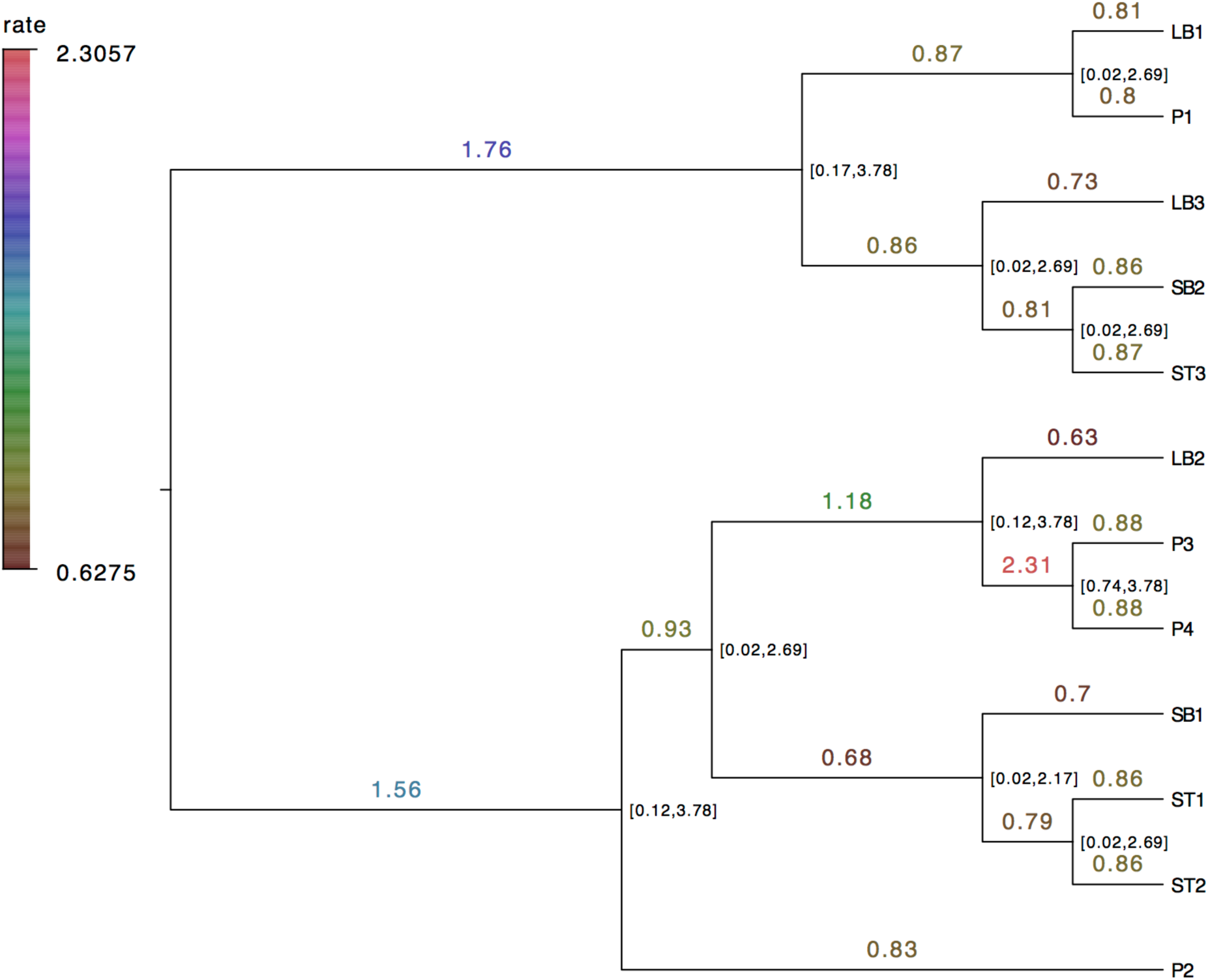
Mean sampled mutation rates using UCED clock model assuming a fixed MP tree for mouse_2. Branch labels show mean mutation rates and node labels provide the 95% HPD interval (see text). Values are rounded to 2 decimal places. The colored bar shows the color codes of the mean rates.

**Supplementary Figure S9:**
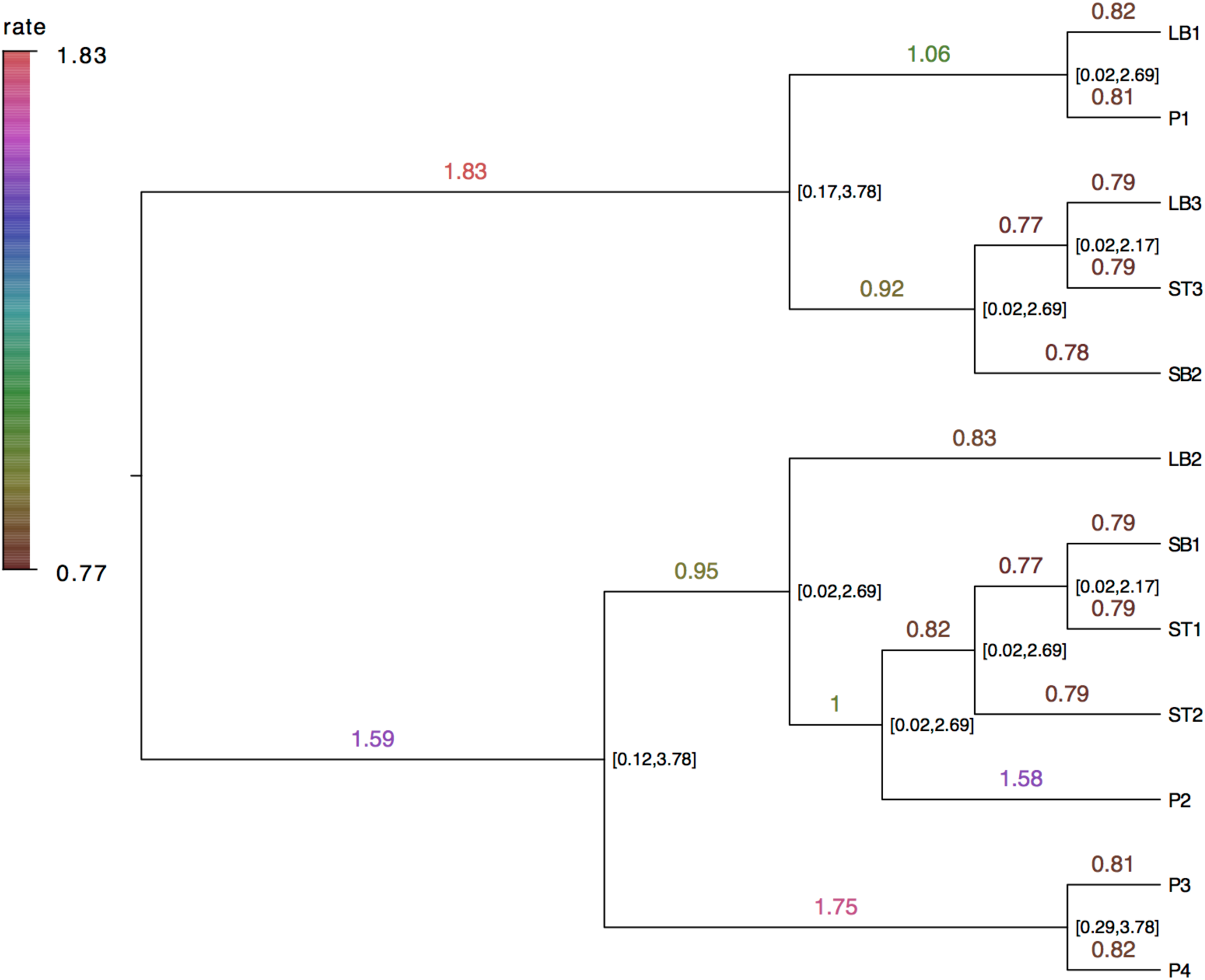
Mean sampled mutation rates using UCED clock model with unconstrained tree topologies for mouse_2. Branch labels show mean mutation rates and node labels provide the 95% HPD interval (see text). Values are rounded to 2 decimal places. The colored bar shows the color codes of the mean rates.

The range of sampled mean rates is considerably bigger for UCED than for UCLD and can be explained with the lognormal being unimodal, and has more of its probability mass around the mode, while the exponential has more probability mass around 0.

### Gamma rate heterogeneity

Substitution rates can be different across different sites within the same sequence and the gamma model has been widely used to fit the data (Yang 2014) and it is also available in BEAST 2. Gamma rate heterogeneity with four different categories was selected with the different clock settings, other parameters remaining the same, and marginal likelihood was estimated with 32 steps and the MCMC chain length set to 300000. The average of two technical replicates were used to calculate the Bayes Factor compared to the corresponding settings without gamma rate heterogeneity added. The models without gamma rate heterogeneity have consistently outperformed the corresponding models with gamma rate heterogeneity added, the lnBF being in the 2.2 - 3.4 range. Please see Supplementary Table S1 below.

**Supplementary Table S1:** Log marginal likelihood estimates for different settings with and without gamma rate variation via path sampling for mouse_1 and mouse_2. Values without gamma rate heterogeneity (mean of three technical replicates) highlighted in blue, values with gamma rate heterogeneity (mean of two technical replicates) added highlighted in bordeaux and calculated log Bayes Factor displayed in purple. Besides the clock models, other models and parameters were set constant: the reversible-jump based model was used for substitution model, coalescent tree prior with constant population was used for the branching process.

| no gamma rate variation<br>with gamma rate variation<br>ln(Bayes Factor) | Strict, 32<br>steps | UCED, 32<br>steps | UCLD, 32<br>steps |
| --- | --- | --- | --- |
| mouse_1<br>fixed MP tree | -119.5<br>-122.9<br>3.4 | -119.5<br>-122.8<br>3.3 | -119.4<br>-122.7<br>3.3 |
| mouse_1<br>unconstrained | -141.3<br>-144.5<br>3.2 | -140.9<br>-144.3<br>3.4 | -141.2<br>-144.4<br>3.2 |
| mouse_2<br>fixed MP tree | -77.1<br>-79.6<br>2.5 | -75.9<br>-78.3<br>2.4 | -76.7<br>-79.1<br>2.4 |
| mouse_2<br>unconstrained | -92.6<br>-95.0<br>2.4 | -91.4<br>-93.9<br>2.5 | -92.3<br>-94.5<br>2.2 |

### Reversible-jump based model for substitution model selection

Instead of using Bayes Factor based model comparison and selection with the numerous substitution model variants available in BEAST 2 we have used the reversible-jump based model (RB) add-on from the RBS package^19^. The RB model automatically converges to a model with a number of parameters supported by the data and jumps from model to model in a hierarchy of models throughout the MCMC run.

Supplementary Table S2 below shows 6 RB traces from the combined logs of the mouse_1 Strict clock with fixed MP tree and mouse_2, UCED relaxed clock with fixed MP tree runs.

**Supplementary Table S2:** RB traces of different substitution models for mouse_1 and mouse_2. The combined logs have been used from 3 different runs. For mouse_1 Strict clock and fixed MP tree were set, for mouse_2, UCED relaxed clock were set under fixed MP tree.

| RB traces | mouse_1, mean<br>Strict, fixed MP | mouse_1<br>ESS | mouse_2, mean<br>UCED, fixed MP | mouse_2<br>ESS |
| --- | --- | --- | --- | --- |
| RBcount | 2.155 | 8179 | 1.531 | 8092 |
| RBrates.1 | 0.39 | 4758 | 0.223 | 3513 |
| RBrates.2 | 3.249 | 12703 | 1.193 | 7375 |
| RBrates.3 | 0.278 | 3967 | 0.203 | 2137 |
| RBrates.4 | 0.126 | 6018 | 0.184 | 1382 |
| RBrates.5 | 0.465 | 1994 | 0.267 | 587 |

Depending on the value of X of RBcount all rates numbered up to X are used, the resolution of the RBRates being: 0 = K81, 1 = HKY85, 2 = TAN93, 3 = TIM, 4 = EVS and 5 = GTR. Rates that are not used will be sampled with low frequency, so even though all rates are reported in the log these will probably show up with a low ESS in Tracer. Looking at the RBcounts, rate mean values and ESS-s in Supplementary Table S2 for both mice the TAN93 model is used predominantly. At this point re-running the samplers with this substitution model would be double dipping leading to the artificial reduction of variability in the data interpretation.

### Tree Prior Selection: coalescent models and Skyline plots

The coalescent models have consistently outperformed the Yule tree prior, or the Birth Death model via Bayes Factor comparison by path sampling. For the MCMC runs for path sampling the unconstrained runs have been used without a fixed tree. After choosing the coalescent branching process the next question was to decide whether a constant population should be assumed or an exponentially growing one. One would be tempted to assume that during mammalian embryogenesis the growth of the cell populations are exponential due to constant mitotic doublings but it should be also considered that apoptosis and necrosis events keep the population more at bay or on a less than exponential curve. Fortunately in BEAST 2 the Bayesian Skyline plot method is available and population history can be reconstructed assuming enough data or at least important hints can be gained on population dynamics.

Supplementary Figure S10 shows two Bayesian Skyline Plots based on unconstrained runs for mouse_1 and mouse_2, respectively. For mouse_1 the Strict clock and RB substitution model was specified. For mouse_2, UCED and RB were given.

**Supplementary Figure S10:**
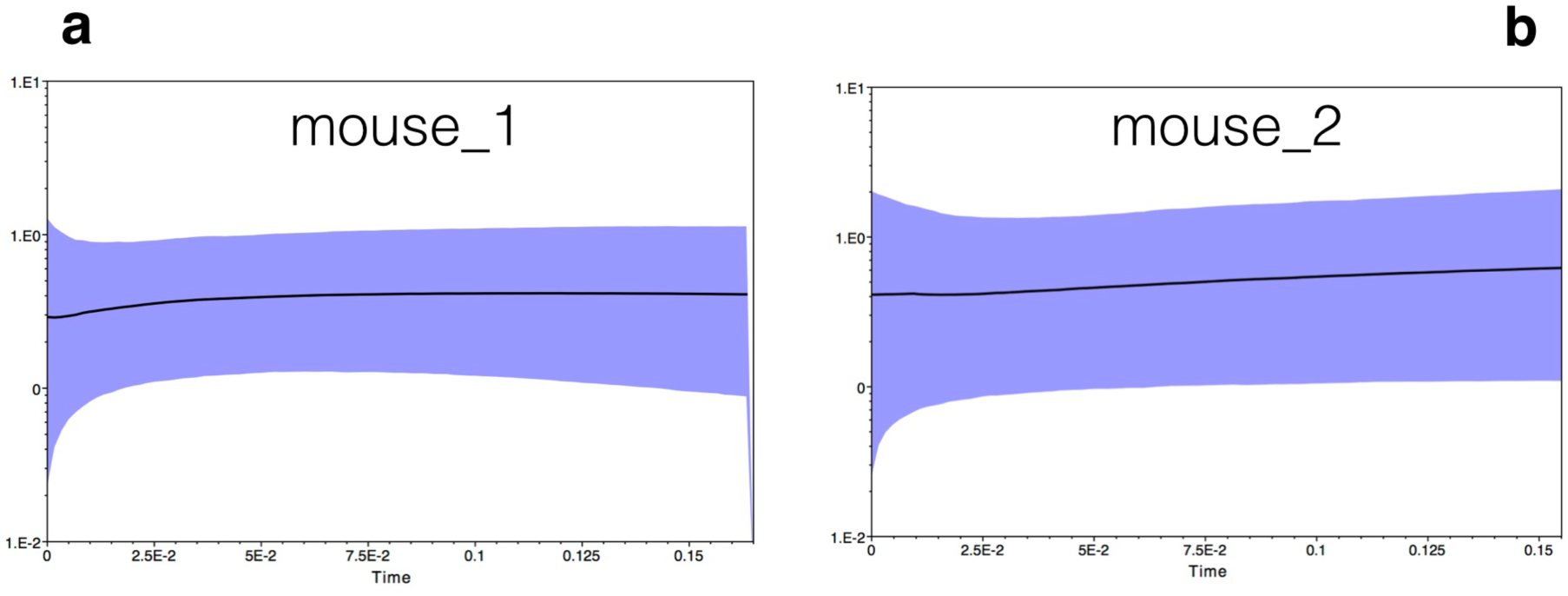
Bayesian Skyline Plots based on unconstrained runs for mouse_1 and mouse_2. **A**., Mouse_1, strict clock and RB substitution model **B**., Mouse_2, UCED clock and RB.

Based on the plots the data is not enough to provide information on population changes throughout so we have used the coalescent prior with the constant population assumption for the runs evaluating the clock models.

